# Mapping human–macaque encounter potential by integrating visitor-use and macaque-use intensity in Kamikochi, Japan

**DOI:** 10.64898/2026.07.30.741901

**Authors:** Yosuke Otani, Ayaka Tsuchihashi, Satoru Sugita, Hiromichi Fukui, Yusuke Shikano, Takuya Matsumoto

## Abstract

Managing visitor–wildlife interactions in high-use protected areas requires identifying not only where visitors concentrate but also where visitor use overlaps with wildlife space use. We estimated relative human– macaque encounter potential in Kamikochi, Chubu Sangaku National Park, Japan, where Japanese macaques (Macaca fuscata) and visitors use the same frontcountry landscape. We integrated Mobile Spatial Statistics derived from mobile phone network data, trail-camera counts, and direct-follow location data from macaques along a frontcountry–backcountry gradient. Broad-scale 500-m grid estimates of daytime visitor presence for the 2023 and 2024 open seasons were allocated to 50-m trail sections and adjusted using counts from 10 trail cameras. Automated video counts based on YOLO11 and ByteTrack showed high agreement with manual counts of 100 validation videos (MAE = 0.10 persons/video, MAPE = 1.26%, exact agreement rate = 92.0%). Visitor-use and macaque-use data were converted into kernel density surfaces, and encounter potential was calculated as their normalized product. High encounter potential was concentrated in the central frontcountry: 87.1% of areas with index values ≥ 0.9 fell within the frontcountry and 12.9% within the backcountry. Areas with high visitor use or high macaque use alone did not necessarily show high encounter potential when the two distributions were spatially offset. These results show that spatial overlap provides management information not captured by visitor counts alone. Relative encounter potential can help prioritize patrols, visitor messaging, viewing management, and other interventions in protected areas where wildlife does not consistently avoid heavily used visitor spaces.

## 1. Introduction

In high-use national parks and nature reserves, a basic management challenge is to maintain visitor experiences while conserving natural resources. In this context, visitor management requires not only estimating visitor numbers but also identifying where visitor use is concentrated and which natural resources or wildlife species may be affected. Especially in protected areas where wildlife and tourists use the same places, human use alone is insufficient for identifying management priorities. Operational decisions such as trail improvement, patrols, warnings, and use regulation therefore require a perspective that evaluates the use of space by both people and wildlife on a common spatial basis (Leung & Marion, 1999).

Many studies of recreation impacts in protected areas have reported that increasing human use can reduce wildlife use near trails, shift activity into the night, or contribute to spatial separation from areas used by people. In Glacier National Park, camera-trap data showed that recreational hiking was associated with reduced trail use and shifts in daily activity patterns in multiple mammal species, including large carnivores and ungulates (Anderson et al., 2023). In Norwegian national parks, overlaying tourist trail use with GPS data from wild reindeer *(Rangifer tarandus)* revealed large-scale segregation between people and animals and led to management implications including visitor-use adjustment (Gundersen et al., 2019). White-tailed deer *(Odocoileus virginianus)* have likewise been reported to reduce their spatiotemporal overlap with hikers in a protected area (Millien et al., 2024). Together, these studies show that avoidance, displacement, and temporal adjustment are important and well-documented responses of wildlife to recreational use, although such responses may vary among taxa, sites, and behavioral contexts.

This body of work also aligns with spatial strategies in protected-area management. Visitor management has long emphasized zoning, concentration of use, dispersal of use, and closure as basic strategies for spatially regulating contact between people and natural resources (Leung & Marion, 1999). Yet implementing such strategies in the field requires data showing where visitors stay and where they simply pass through. In recent years, location data derived from mobile devices have been widely used for visitor monitoring and have proved useful for estimating visitation patterns and spatial density in protected areas (Creany et al., 2021; Kim et al., 2023). However, such data are affected by sampling biases and limits in spatial resolution and are not, by themselves, sufficient for segment-level management decisions (Miyasaka et al., 2018).

Research on wildlife space use has become increasingly detailed through advances in GPS telemetry and camera trapping. At the same time, a global systematic review of non-consumptive recreation reported that documented effects on animals include behavioral responses such as increased flight and vigilance, changes in spatial or temporal habitat use, and declines in abundance, occupancy, or density (Larson et al., 2016). For wild primates, however, responses to people may also be shaped by prior exposure, habituation, provisioning history, and site-specific socioecological contexts, and therefore are not always well characterized as simple spatial avoidance. In primate field studies, habituation without provisioning has long been used to enable close-range observation of wild groups (Williamson & Feistner, 2003; Hernández Tienda et al., 2022). Japanese macaques *(Macaca fuscata)* on Yakushima provide a relevant example: dietary and ecological studies of non-provisioned Yakushima macaques have shown that their foraging is based on naturally occurring forest resources and varies with seasonal food availability (Hanya, 2004; Agetsuma & Nakagawa, 1998), yet visitors can still encounter and observe macaques at close range along roads in the island’s forested landscape (Yakushima Town, n.d.). This indicates that close-range primate viewing can arise not only from tourist feeding or formal provisioning, but also from the spatial overlap of human travel routes, habituated or human-tolerant macaques, and natural foraging habitats. These examples suggest that, for wild primates, visitor management should consider context-dependent patterns of space use and human tolerance rather than relying solely on the expectation that wildlife will spatially separate from areas of human use.

Kamikochi, part of Chubu Sangaku National Park, illustrates why this perspective is relevant. The central walking area contains hotels and major tourism nodes and is served by flat, accessible trails. At the same time, river terraces, grasslands, and waterside environments provide valuable habitat and foraging opportunities for Japanese macaques (Milner et al., 2021; Nagahara et al., 2025). The frontcountry may therefore function as a shared-use landscape that is attractive to both visitors and macaques. In Japan, feeding wild mammals and birds in national parks is regulated, and Kamikochi explicitly publicizes a no-feeding rule (Ministry of the Environment, Japan, 2021; Kamikochi Official Website, n.d.). Nevertheless, Kamikochi provides opportunities for visitors to observe Japanese macaques at close range, and the degree of proximity between visitors and macaques can itself become a management concern.

Against this background, we integrated mobile-based estimates of visitor use, trail-camera-based visitor counts, and Japanese macaque location data to estimate encounter potential as a relative index of spatial overlap between visitor use and macaque space use. We evaluated whether this index could help identify sections where patrols, visitor warnings, and other management efforts may be prioritized along a frontcountry– backcountry gradient. The first contribution of this study is to visualize where management resources may be most effectively allocated within the study area, thereby providing spatial evidence relevant to visitor-management practice. Second, by treating human–wildlife spatial overlap as a management indicator that cannot be inferred from visitor counts alone, the study links recreation-impact research with section-level management in protected areas. More broadly, it highlights the need to adapt management of high-use frontcountry areas to situations in which wildlife, such as primates, continues to use spaces intensively used by visitors rather than consistently avoiding them.

## 2. Methods

### 2.1. Study area and zoning

Kamikochi, the study site, is part of Chubu Sangaku National Park and is a mountain scenic area located at an elevation of approximately 1,500 m. It is a high-use protected area with concentrations of hotels, tourism facilities, and flat, well-maintained trails, and receives approximately 1.3–1.5 million visitors annually (Kamikochi Liaison Council of Chubu Sangaku National Park, 2026). Kamikochi is officially open each year from April 17 to November 15. During the closed season, heavy snow accumulation makes regular tourism use difficult; transportation services such as buses and taxis are suspended, and visitor services such as accommodation and food service are not provided. Accordingly, tourism use is concentrated during the open season.

For zoning, we used the Kamikochi zoning map adopted by the Kamikochi Liaison Council of Chubu Sangaku National Park. In this study, we treated the strolling area as the frontcountry and the nature appreciation area as the backcountry. Figure 1 shows the study area and zoning categories. The frontcountry contains the main visitor-use nodes, including hotel clusters, the visitor center, and the bus terminal. Most trails in this zone are barrier-free and accessible to wheelchair users. By contrast, the backcountry provides a more natural recreation setting, and trekking shoes are recommended for these trails. Kamikochi also includes trekking and mountaineering zones, which are intended for visitors engaged in more demanding hiking and mountaineering. These zones were excluded from the present study because they are outside the main frontcountry–backcountry gradient examined here.

**Figure 1.**
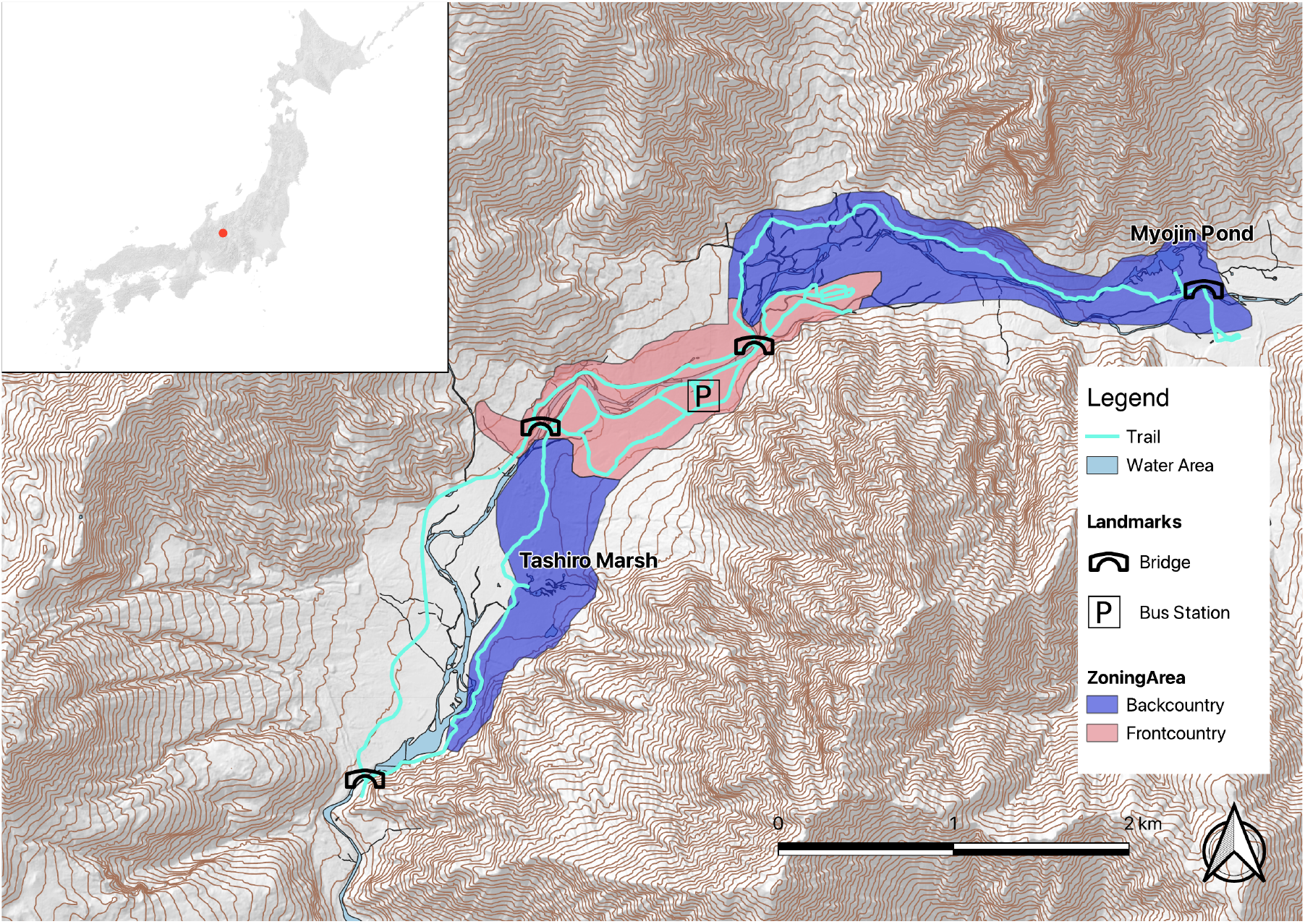
Study area and zoning categories in Kamikochi, Japan. Caption: The map shows the study area in Kamikochi, Japan, including the trail network and the frontcountry and backcountry zones used in this study. The inset indicates the location of Kamikochi within Japan. Topography and contour lines were derived from the 10-m digital elevation model of the Geospatial Information Authority of Japan, and zoning boundaries were prepared by the authors based on materials from the Ministry of the Environment, Japan.

### 2.2. Spatial datasets

We integrated the Kamikochi zoning map, trails, 500-m grid population data, trail-camera counts, and Japanese macaque location data. The study extent ran from the beginning of the main walking trail in Kamikochi (southwestern part of Fig. 1) to the zoning boundary between the strolling area (frontcountry) and the nature appreciation area (backcountry) (northeastern part of Fig. 1). The trail line within the study extent was prepared in GIS using satellite imagery and maps from the Geospatial Information Authority of Japan and was corrected by overlaying track logs recorded by the author while walking the trails with a handheld GPS device. Trail sections that were closed to visitors during the study period because of slope failures or other hazards were excluded from the trail network used in the analysis. The total trail length within the study extent was 17,504 m. This trail network was divided, in principle, into 50-m sections, which served as the basic unit for the human-use analysis described below. Because some sections were shorter than 50 m at trail intersections and similar locations, the total number of trail sections was 360. Spatial processing and mapping were conducted in QGIS 3.40 (QGIS Development Team, 2024), and some numerical processing was performed in Python 3.12 (Python Software Foundation, 2023).

### 2.3. Large-scale visitor-use data from Mobile Spatial Statistics

To characterize broad-scale visitor use in Kamikochi, we used Mobile Spatial Statistics (MSS) provided by DOCOMO InsightMarketing, Inc., based on NTT DOCOMO’s mobile phone operational data (DOCOMO InsightMarketing, n.d.). MSS are population statistics estimated from mobile phone operational data after privacy-preserving processing, including anonymization, aggregation, and confidentiality processing, and do not identify individual users. The dataset used in this study was the population distribution dataset for 28 cells of the 500-m grid covering the study area (Fig. 2). It provided estimated population values for each grid cell, date, and hourly time interval.

**Figure 2.**
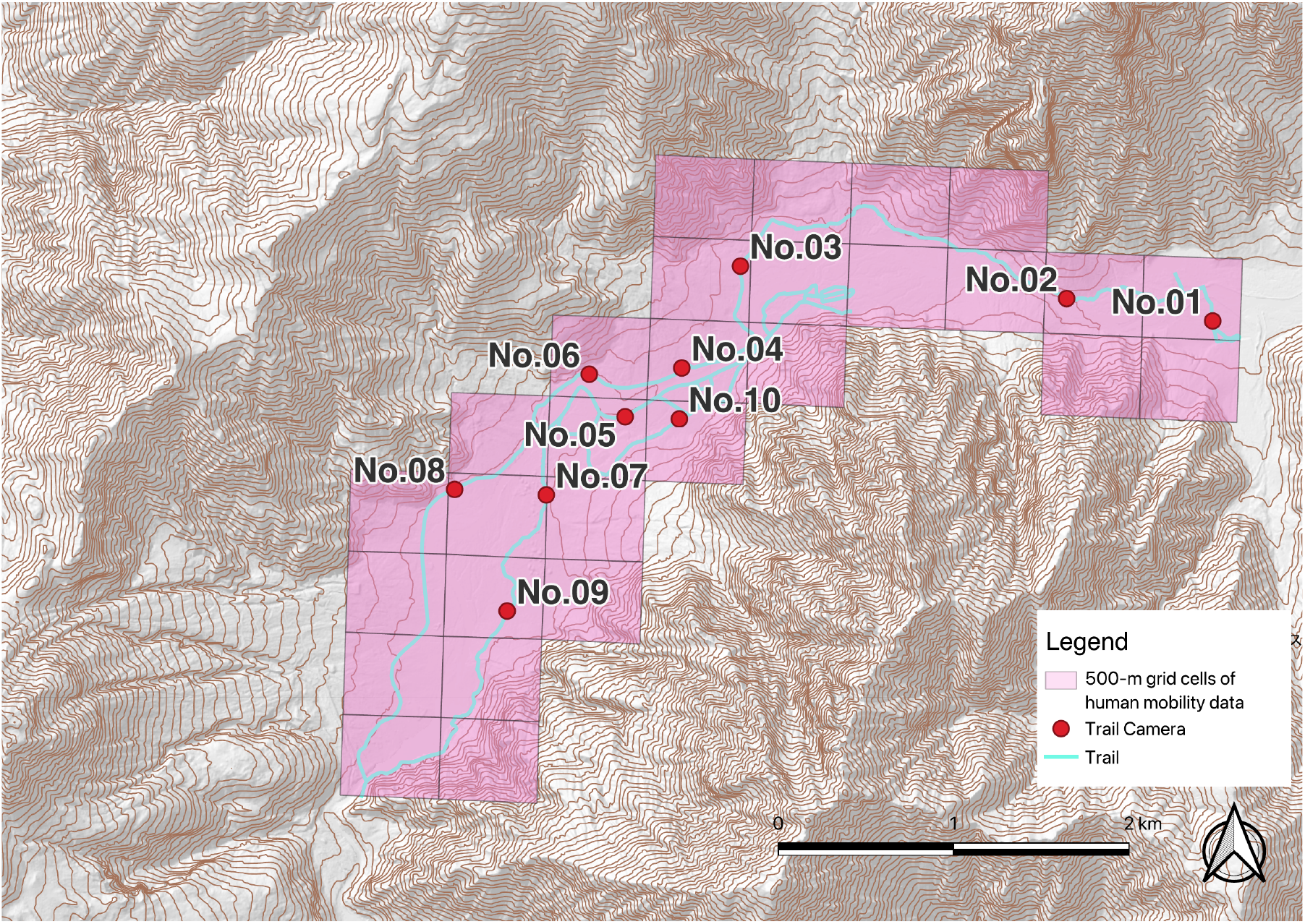
500-m grid cells of human mobility data and trail camera locations. Caption: Pink polygons indicate the 500-m grid cells used for human mobility data, red points indicate trail camera locations, and blue lines indicate trails.

The study period covered the full open season of Kamikochi in two years: April 17–November 15, 2023, and April 17–November 15, 2024, corresponding to 426 days in total. For each grid cell, we used only daytime hourly intervals from 06:00 to 18:00, corresponding to the period from 06:00 to 18:59, and summed the estimated population values across these time intervals and dates to obtain a cumulative person-hour index of visitor presence. Thus, MSS values were treated as broad-scale indicators of daytime human presence rather than as direct counts of trail users. Although MSS can capture long-term and spatially consistent patterns of human use, its 500-m spatial resolution is too coarse to represent the detailed trail network of Kamikochi. In some cases, a single grid cell contains multiple trail sections as well as both riverbanks. We therefore downscaled and corrected the grid-level MSS data to the trail-section level using local visitor counts from trail cameras, as described below.

### 2.4. Trail-camera survey of visitor passage

To locally calibrate the visitor-use data from MSS, we installed 10 trail cameras (400-CAM092, SANWA SUPPLY) along trails in the study area and recorded videos of passing park visitors (Fig. 2). Four cameras were placed in the frontcountry and six in the backcountry. To minimize impacts on protected forest and other natural features, installation sites were selected after field inspections conducted together with Ministry of the Environment staff.

Videos were recorded from October 2 to October 29, 2024, and each clip was saved as a 10-s video. To reduce the likelihood of identifying individuals, cameras were positioned low and aimed at visitors’ feet and lower bodies. All video data were handled in accordance with personal-information protocols, stored on an encrypted hard disk drive, and kept in a locked cabinet. All analyses were conducted in a local computing environment.

### 2.5. Counting people in videos and validating automated counts

The number of people passing in each video was estimated by combining the object-detection model YOLO11 (Jocher & Qiu, 2024) with the multi-object tracking method ByteTrack (Zhang et al., 2022). We used the weight file yolo11n.pt and ran inference on a CUDA-enabled GPU (RTX4000Ada, NVIDIA). Videos recorded at 30 fps were read frame by frame, and inference was run once every three frames. Only the person class was detected. Detections were passed to ByteTrack, track IDs were assigned, and the number of passing visitors was summarized from track information for each video.

The main ByteTrack settings were track_thresh = 0.5, track_buffer = 30, match_thresh = 0.8, aspect_ratio_thresh = 1.6, min_box_area = 10, mot20 = False, and frame_rate = 30. To validate the automatic counts, 100 videos were randomly sampled from the full set of 30,339 videos and compared with manual visual counts. We compared AI-based estimates and manual counts at the video level and assessed mean absolute error (MAE), mean absolute percentage error (MAPE), exact agreement rate, and the typical causes of counting error.

### 2.6. Downscaling visitor-use data and deriving the relative visitor-use indicator

We downscaled the grid-level visitor-use data from MSS to 50-m trail sections and adjusted the resulting values using trail-camera counts. This procedure was designed to retain the broad spatial coverage of mobile-phone-based population data while translating it into a spatial resolution suitable for section-level visitor management.

The base visitor-use value for trail section *s*, *P(s)*, was obtained by allocating the cumulative person-hour index in grid cell *m*, *P(m)*, in proportion to the length of section *s*, *ℓ(s)*, relative to the total trail length within the same grid cell, *L(m)*:

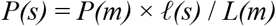

For each camera *c*, we calculated a relative camera weight, *K(c)*, as the proportion of visitors recorded at that camera, *n(c)*, relative to the total number of visitors recorded across all cameras:

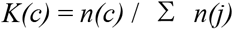

where the summation is over all cameras *j*.

The adjusted relative visitor-use indicator for section *s*, *P*(s)*, was then calculated as:

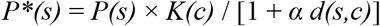

where *d(s,c)* is the distance in meters between section *s* and the corresponding camera *c*, and *α* = 0.01 m⁻¹ is the distance-decay parameter. For each 50-m trail section, the relative camera weight of the nearest camera on the same route was assigned. The resulting *P*(s)* values integrate broad-scale estimates of daytime visitor presence, local observations of visitor passage, and a distance-decay effect from the assigned trail camera. These values should therefore be interpreted as relative indicators of visitor use, rather than as estimates of absolute visitor numbers.

### 2.7. Japanese macaque location data

Japanese macaque space use was characterized by directly following three groups that ranged into the study area: the KM, KK, and KT groups. The procedure used to locate macaque groups before direct follows is described in Tsuchihashi et al. (2025). Locations were recorded every 30 s using a handheld GPS receiver. Field surveys were conducted from May 11, 2023, to November 17, 2024, and only data collected during the Kamikochi open seasons were extracted for analysis. The final dataset comprised 50 survey days, totaling 88 h of direct-follow data.

These data were collected mainly on or near trails. Because our objective was to evaluate spatial overlap in land use between visitors and macaques, the key information required was where macaques used areas that overlapped with visitor travel spaces within the focal management area. Accordingly, we treated these data not as a complete record of macaque movements across all of Kamikochi, but as a dataset focused on macaque use in areas where overlap with visitor use was most relevant. To visualize broad patterns of macaque use near trails, macaque locations were converted into a kernel density surface using a bandwidth of *h*=500 m.

### 2.8. Definition and normalization of encounter potential

In this study, we defined encounter potential as a relative index of spatial overlap between visitor use and macaque space use. Specifically, encounter potential was calculated as the product of the visitor-use intensity surface and the macaque-use intensity surface. To make the two surfaces comparable in format, we first assigned the adjusted relative visitor-use indicator *P*(s)* to the centroid of each 50-m trail section. These values were then converted into a continuous visitor-use intensity surface using kernel density estimation with a bandwidth of *h* = 50 m. For visitor use, this bandwidth was chosen as a local smoothing parameter that matched the scale of the 50-m trail sections and avoided excessive smoothing across separate trail segments. Because visitor movement in Kamikochi is largely concentrated on trails, this local smoothing approach was considered appropriate for representing trail-based visitor use.

The raw encounter-potential index at location x was calculated as:

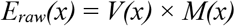

where *V(x)* is the visitor-use intensity at location *x* and *M(x)* is the macaque-use intensity at the same location. For visualization and comparison among sections, this raw index was normalized by the maximum value across the full study extent:

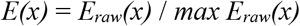

where the maximum value was calculated across the full study extent.

The resulting *E(x)* should be interpreted as a relative index of encounter potential, not as an absolute probability of encounter, conflict, or damage. Rather, it indicates, in relative terms, areas where visitor use and macaque space use overlap strongly and where management attention may need to be prioritized.

### 2.9. Ethics

This study was approved by the Research Ethics Committee of the Center for the Study of Co* Design, The University of Osaka (approval no. 2024-2). All fieldwork complied with relevant laws and regulations, and the required permits were obtained from the Ministry of the Environment, the Forestry Agency, Nagano Prefecture, and Matsumoto City.

## 3. Results

### 3.1. Overview of Mobile Spatial Statistics and trail-camera data

Figure 3 shows the monthly distribution of mean daily estimated visitor population per grid cell based on MSS. Estimated visitor population across Kamikochi showed clear seasonal variation throughout the open season, increasing from spring into summer, reaching high values in August, and then declining toward autumn. This seasonal pattern was broadly similar in 2023 and 2024. At the same time, the boxplots show large differences among grid cells within the same month, indicating strong spatial heterogeneity in where visitors were present. These results indicate that management needs to consider not only monthly variation in overall use but also the spatial structure of visitor-use concentration.

**Figure 3.**
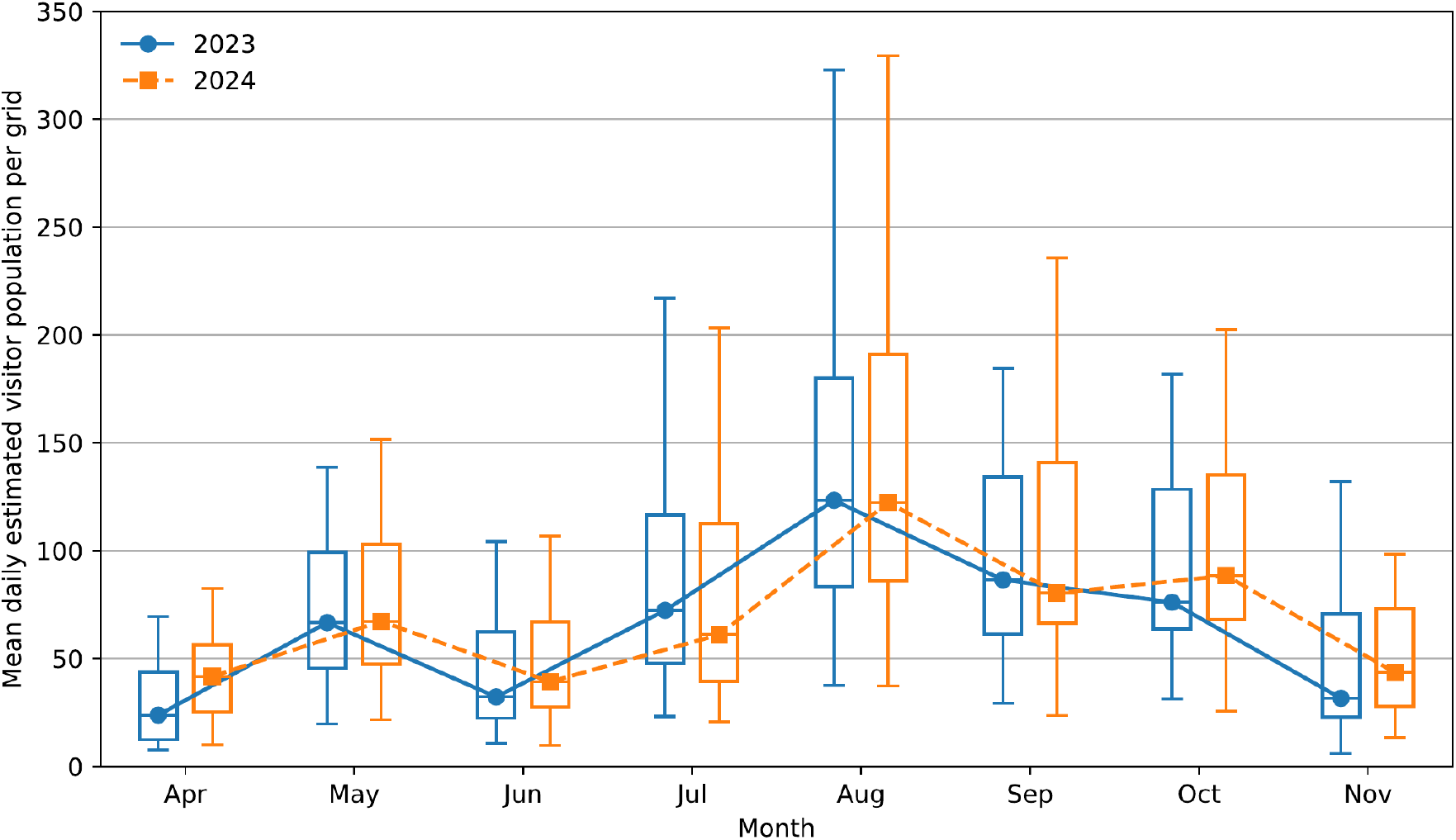
Monthly variation in mean daily estimated visitor population per grid in 2023 and 2024. Caption: Boxplots show the distribution of mean daily estimated visitor population per grid within each month, based on mobile spatial statistics data in 2023 and 2024. Lines connect the monthly medians for each year.

Table 1 summarizes the deployment period of each trail camera, the number of video clips, the number of people recorded, and the mean number of people recorded per effective operating hour. Across the 10 cameras, a total of 30,339 video clips were obtained, and 93,872 people were recorded. The mean number of people recorded ranged from 4.4 to 191.8 people/h, showing large differences among sites. For example, cameras 5 and 6 were located close to one another (Fig. 2), yet their mean counts were 56.7 and 96.6 people/h, respectively. This indicates that visitor passage and concentration along the trail system were not spatially uniform and were not explained by distance alone.

**Table 1.** Deployment period, video volume, and visitor counts recorded by trail cameras.

| Camera No. | Start date | End date | Number of videos | Number of people recorded | Mean number of people recorded per hour |
| --- | --- | --- | --- | --- | --- |
| No.01 | 2024-10-03 | 2024-10-30 | 3,186 | 9,895 | 129.5 |
| No.02 | 2024-10-03 | 2024-10-30 | 3,194 | 9,450 | 128.6 |
| No.03 | 2024-10-03 | 2024-10-30 | 3,058 | 10,432 | 191.8 |
| No.04 | 2024-10-04 | 2024-10-29 | 2,931 | 10,213 | 110.7 |
| No.05 | 2024-10-02 | 2024-10-30 | 3,018 | 7,979 | 56.7 |
| No.06 | 2024-10-02 | 2024-10-29 | 3,068 | 9,275 | 96.6 |
| No.07 | 2024-10-02 | 2024-10-29 | 3,558 | 14,611 | 129.4 |
| No.08 | 2024-10-02 | 2024-10-29 | 1,565 | 2,851 | 4.4 |
| No.09 | 2024-10-02 | 2024-10-29 | 3,317 | 10,676 | 113.3 |
| No.10 | 2024-10-02 | 2024-10-30 | 3,444 | 8,490 | 21.0 |
| **Total** |  |  | **30,339** | **93,872** | **52.4** |

Note. Start and end dates indicate the deployment period of each trail camera. Number of videos indicates the total number of video files obtained from each camera. Mean number of people recorded per hour was calculated from the number of people recorded and the effective operating time of each camera.

We counted people in the trail-camera videos using the AI-based procedure described above. To evaluate the validity of this automatic counting procedure, 100 videos randomly sampled from the 30,339 clips were compared with manual visual counts. The mean absolute error was 0.10 persons/video, the mean absolute percentage error was 1.26%, and the exact agreement rate was 92.0%. In 99 of the 100 videos, the difference between AI counts and manual counts was within ±1 person. The total AI count for the validation subset was 315, compared with a manual total of 313, indicating only slight overall overcounting.

### 3.2. Distribution of park visitors along trails

Using the procedures described in Methods 2.6, the cumulative MSS person-hour index in each grid cell was allocated to 50-m trail sections, adjusted using observed trail-camera counts, and converted into the adjusted relative visitor-use indicator *P*(s)*. Using the procedure described in Methods 2.8, we then generated a continuous visitor-use intensity surface along the trail system (Fig. 4). The resulting distribution of park visitors along trails showed clear spatial heterogeneity. The highest values were concentrated in the central part of the study area (Fig. 4: 3-c, d; 4-c, d), largely overlapping the strolling area in the Kamikochi zoning system, i.e., the frontcountry. Relatively high-use areas were also detected around Myojin Pond in the eastern part of the study area (Fig. 4: 2-i, 3-i) and around Tashiro Marsh in the western part (Fig. 4: 6-b), but the sections linking these areas showed relatively low values. The Myojin sector includes a shrine, a restaurant, accommodation, and viewpoints of Mt. Myojin, whereas the Tashiro sector includes marshland and viewpoints. Thus, visitor use was not distributed evenly across the entire trail network but was concentrated at major attraction nodes and lower along the sections used mainly for movement between them.

**Figure 4.**
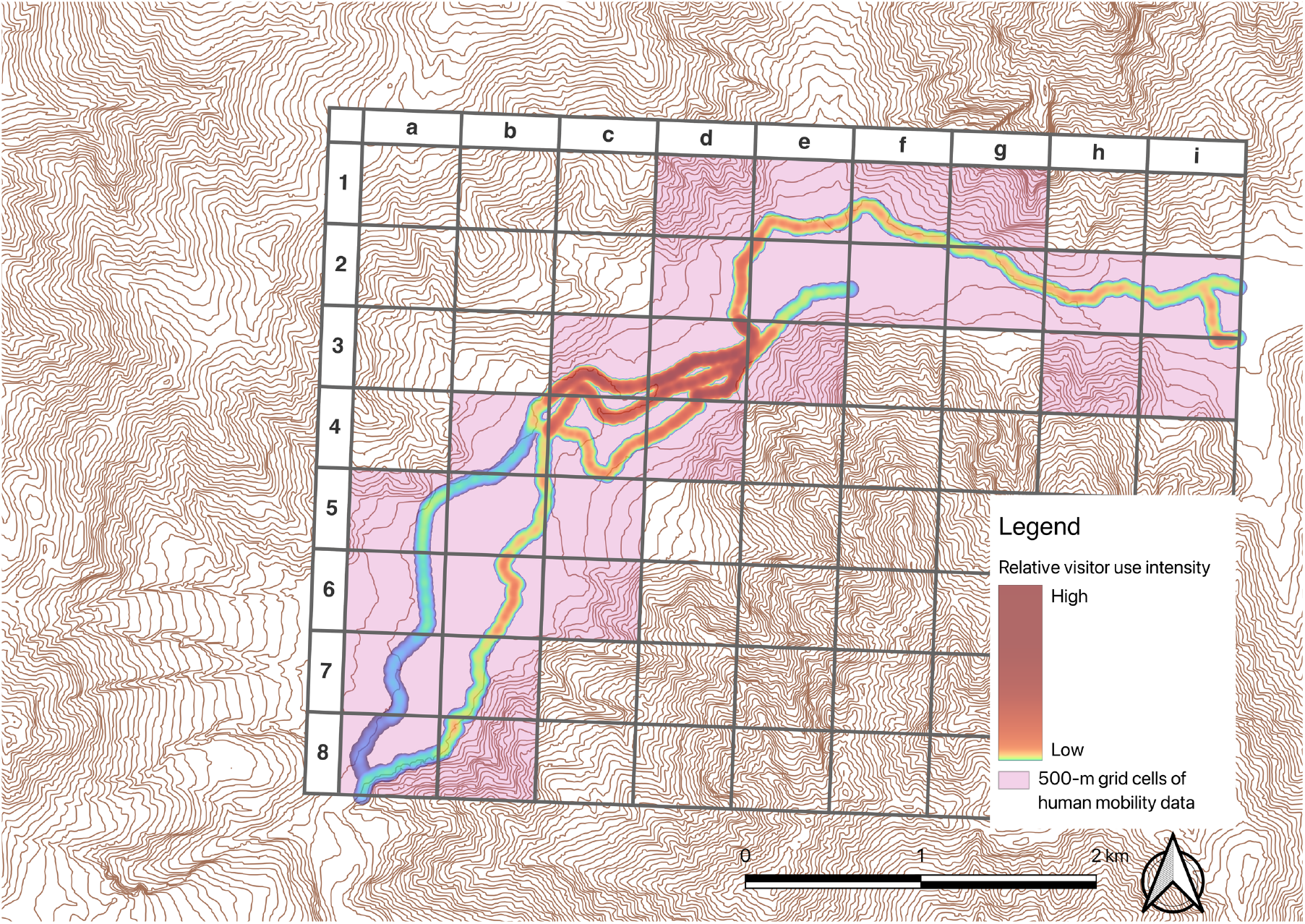
Spatial pattern of relative visitor use intensity along trails in Kamikochi. Caption: The map shows the spatial pattern of relative visitor use intensity along trails in Kamikochi. Visitor population derived from mobile spatial statistics was distributed to 50-m trail segments, corrected with trail camera observations, and smoothed using kernel density estimation to generate a continuous use-intensity surface. Warmer colors indicate higher relative visitor use intensity. Pink polygons indicate the 500-m grid cells for which human mobility data were available, and the alphanumeric grid labels are provided to facilitate reference to specific locations in the text.

### 3.3. Spatial use by Japanese macaques

Kernel density estimation based on location data from the three groups, KM, KK, and KT, showed that Japanese macaque use intensity was highest in the central part of the study area (Fig. 5). Additional areas of moderately high use were detected in the eastern part of the study area, indicating that macaque use was not spatially uniform but included multiple concentration zones. At least within the visitor travel spaces included in the frontcountry and backcountry, macaques therefore showed several areas of concentrated use.

**Figure 5.**
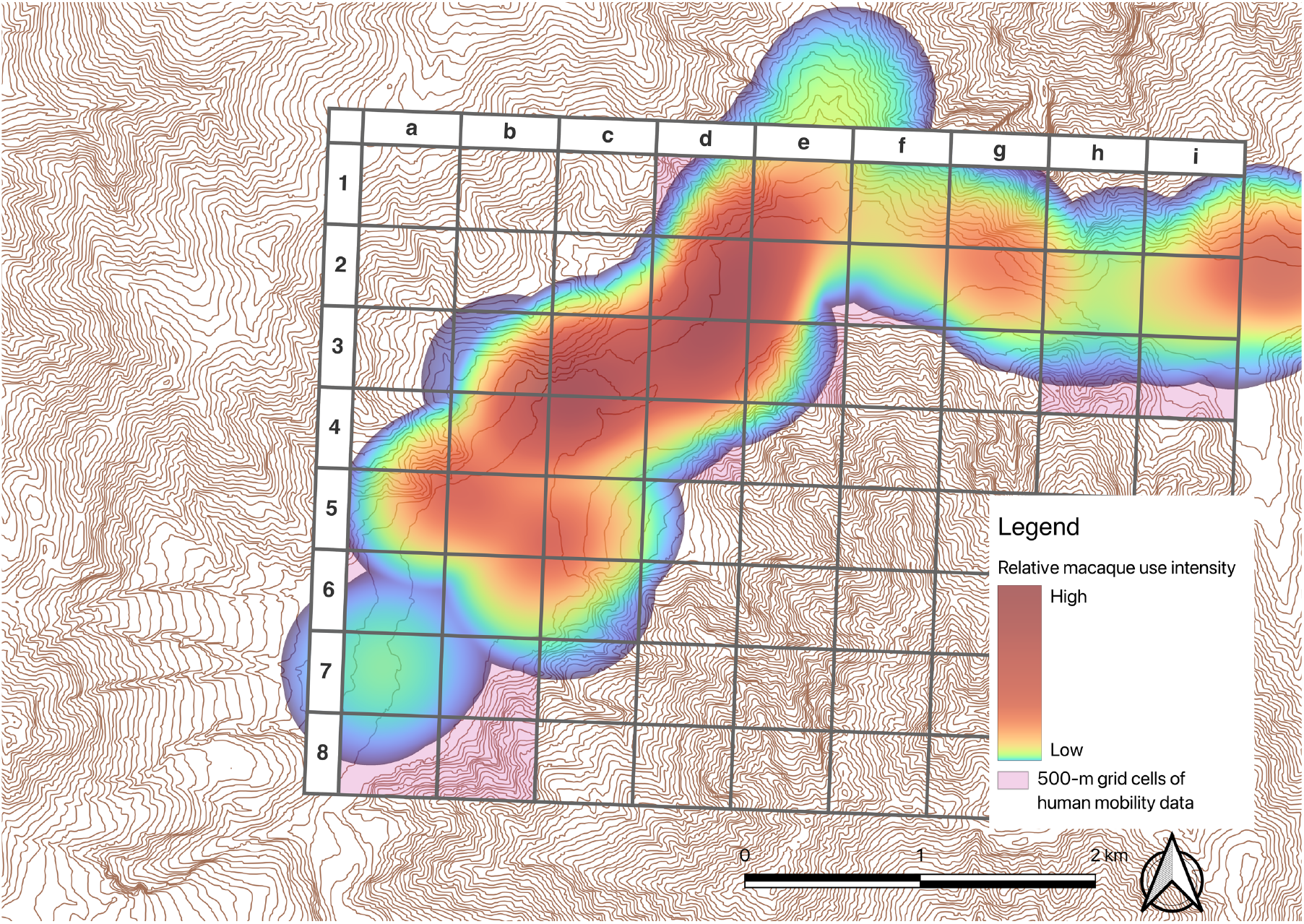
Relative macaque use intensity around trails in Kamikochi. Caption: The map shows the relative macaque use intensity estimated from kernel density analysis of location data from the KM, KK, and KT groups. Warmer colors indicate higher relative macaque use intensity. Pink polygons show the 500-m grid cells with human mobility data, and alphanumeric labels are used to identify specific locations in the text. Because the location data were collected mainly on or near trails, the figure represents macaque use around visitor travel space rather than movements across the entire Kamikochi area.

Because the location data were collected mainly on or near trails, the distribution shown here does not represent macaque movements across the whole of Kamikochi. Rather, it represents patterns of macaque use around visitor travel spaces that are important for evaluating overlap with visitor use. Japanese macaques used both the frontcountry and the backcountry, but areas of macaque use that overlapped most strongly with the trail system were concentrated mainly in the frontcountry.

### 3.4. Spatial pattern of encounter potential and high-risk sections

We multiplied the visitor-use intensity surface by the macaque-use intensity surface and normalized the result to a 0–1 relative index to calculate encounter potential (Fig. 6). In this study, areas with encounter potential values of 0.9 or higher were treated as high-risk areas. Of these high-risk areas, 87.1% fell within the frontcountry and 12.9% within the backcountry. No high-risk areas overlapped with zones outside these two categories. Thus, high-risk areas were not evenly distributed across the park but were concentrated mainly in the frontcountry. By contrast, around Myojin Pond in the east (Fig. 4: 2-i, 3-i) and Tashiro Marsh in the west (Fig. 4: 6-b), both visitor use and macaque use showed relatively high values, but encounter potential was lower than in the central area because the locations of high use by visitors and macaques were slightly offset from one another.

**Figure 6.**
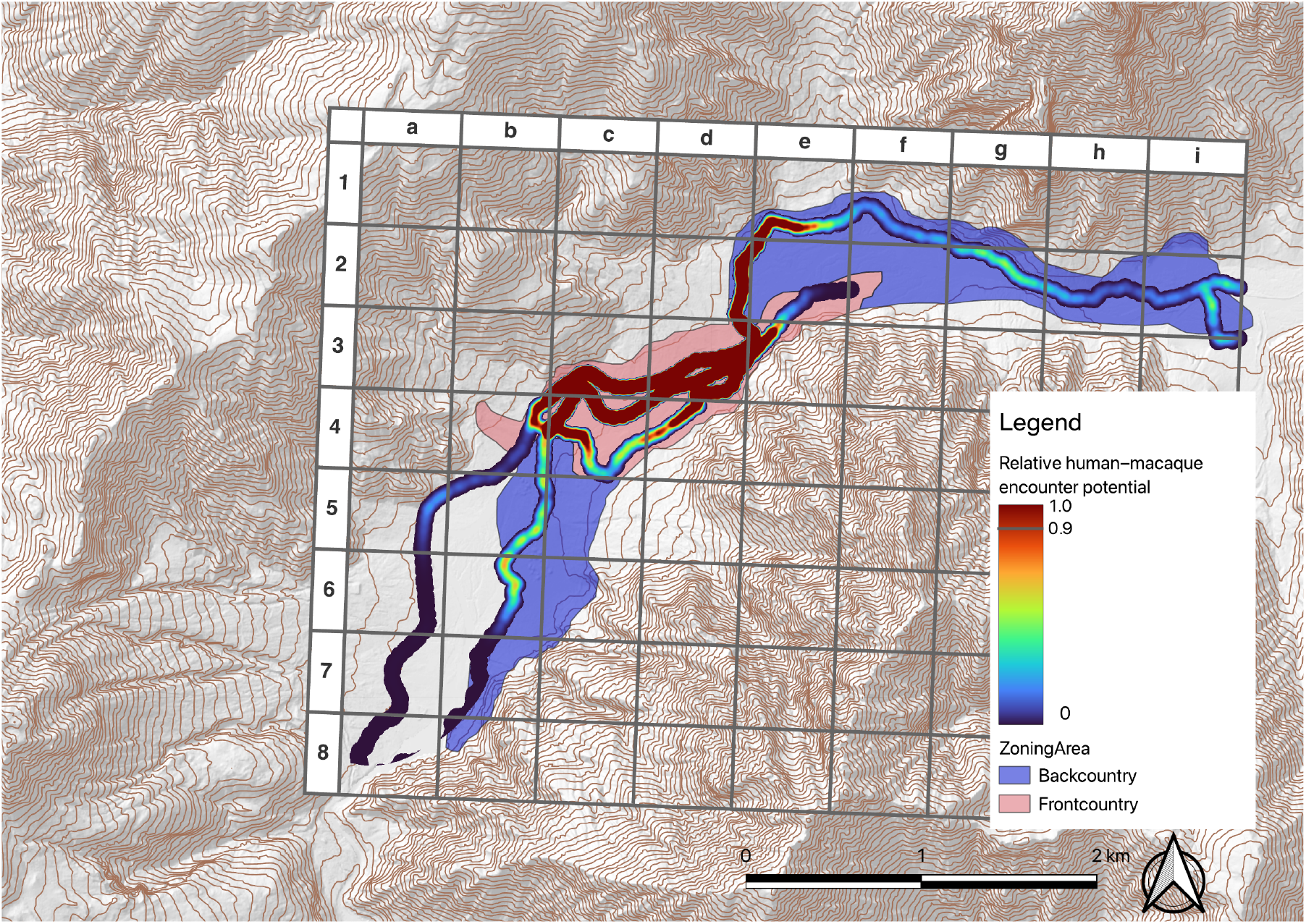
Relative human–macaque encounter potential along trails in Kamikochi. Caption: The map shows the relative human–macaque encounter potential along trails in Kamikochi. Encounter potential was derived by multiplying the visitor and macaque use intensity surfaces and normalizing the result to a 0–1 relative index. Warmer colors indicate higher encounter potential, and values of 0.9 or higher are treated as high-risk sections. Frontcountry and backcountry zones are shown as background polygons, and alphanumeric grid labels are provided to facilitate reference to specific locations in the text.

## 4. Discussion

### 4.1. A shared-use landscape in Kamikochi and implications for visitor management

By integrating broad-scale human mobility data, trail-camera observations of visitor passage, and Japanese macaque location data, this study visualized human–macaque encounter potential in Kamikochi as a relative index. High-risk areas were concentrated in the central frontcountry, corresponding to the strolling area in the zoning system, whereas the backcountry generally showed lower values. This indicates that potential encounters between visitors and macaques are not distributed uniformly across the park but are concentrated in a limited part of the existing frontcountry.

A key result is that high visitor use and high macaque use did not coincide in a simple one-to-one manner. Around Myojin Pond and Tashiro Marsh, both visitors and macaques showed relatively high use, yet encounter potential remained lower than in the central area because their high-use locations were slightly offset. Similarly, trail sections connecting major visitor-use nodes may experience substantial through-traffic, but their encounter potential remained relatively low, probably because visitors pass through these sections with limited dwell time. The indicator developed here is therefore useful not only for identifying areas with many visitors or many macaques separately, but also for detecting management-priority areas that would not be apparent from either distribution alone. Because visitor management in protected areas is fundamentally spatial, involving decisions about where to separate, concentrate use, or prioritize conservation (Leung & Marion, 1999), our results provide practical evidence for such spatial decisions.

The pattern observed in Kamikochi also differs from the commonly reported pattern in recreation-impact studies, in which wildlife reduces use of areas with high human activity or shifts activity in space or time. Studies from Glacier National Park and Norwegian national parks have reported spatial avoidance or temporal shifts in wildlife under increasing human use (Anderson et al., 2023; Gundersen et al., 2019). In Kamikochi, however, visitor use and macaque use still overlapped substantially in the frontcountry. This suggests that the Kamikochi frontcountry functions as a shared-use landscape that is valuable to both visitors and Japanese macaques. In such a setting, the management issue is not only whether people and wildlife can be spatially separated, but also where management attention should be concentrated within shared space. In this sense, the study connects recreation-impact research with section-level visitor management in protected areas.

In Kamikochi, a key management challenge is how to allocate limited resources for patrols, visitor warnings, management of wildlife viewing, and active deterrence. Management attention is required not only for Japanese macaques but also for Japanese black bears *(Ursus thibetanus japonicus)*, sika deer *(Cervus nippon)*, invasive plants, and other concerns (Kamikochi Liaison Council of Chubu Sangaku National Park, 2026). Visitors move across a wide area, and mountaineers may travel early in the morning or late in the day, making it difficult to address all issues uniformly across space and time. Because human and financial resources are limited, and intensive intervention across the entire park is unrealistic in a scenic protected area, managers need spatial evidence to determine where to act first. The encounter-potential index proposed here can support such prioritization and may also be applicable to other mammals that overlap spatially with visitor use.

### 4.2. Implications for non-provisioning primate watching and ecotourism

The significance of this study extends beyond local visitor management in Kamikochi. In some primate tourism settings, wild primates do not fully avoid people but instead share spaces with visitors. At macaque tourism sites in Asia and Barbary macaque tourism sites in North Africa, observation opportunities have often been linked to provisioning, tourist feeding, or dependence on anthropogenic food. In Indonesia, provisioning alters the activity budgets of long-tailed macaques (Ilham et al., 2018), and in Malaysia, human activity shapes macaque behavior at an urban tourism site (Entezami et al., 2024). In Morocco, tourist–macaque interactions frequently involve feeding, and tourist provisioning has been linked to both close contact and health-related concerns in wild Barbary macaques (Maréchal et al., 2016a, 2016b). Recent discussions have therefore emphasized the need to move away from feeding-dependent primate tourism through no-feeding rules, staff intervention, and management of contact opportunities (Entezami et al., 2024; Maréchal et al., 2016a).

The present study shows that opportunities for primate observation are not determined by provisioning alone. In Kamikochi, areas of high encounter potential existed despite the absence of routine provisioning. This finding has both positive and negative implications. On the one hand, opportunities for wild primate watching may be maintained without provisioning when local environmental conditions allow repeated observation. On the other hand, primates that tolerate people may approach visitors at distances close enough to raise management concerns, even in the absence of feeding. In such settings, balancing conservation and tourism requires targeted management based on identifying where visitor use and primate space use overlap.

Kamikochi therefore provides a useful case for considering non-provisioning primate tourism. Where feeding is not used to maintain observation opportunities, managers may first need to identify where encounter potential is concentrated and then focus viewing rules, visitor guidance, ranger deployment, and warning messages in those areas. Thus, the contribution of this study is not limited to Kamikochi; it also offers a practical perspective on spatial prioritization for ecotourism and wildlife-watching systems that aim to avoid provisioning.

### 4.3. Limitations and future directions

This study has several limitations. First, encounter potential is a relative indicator derived from the product of visitor-use intensity and macaque-use intensity; it does not directly estimate the absolute probability of encounters, bites, chasing, or facility damage. Second, macaque location data were collected mainly on or near trails and were not intended to represent complete ranging patterns across the entire Kamikochi landscape. Third, the integrated datasets differed in spatial and temporal resolution. Accordingly, the main strength of this study lies not in estimating absolute levels of risk, but in identifying relative priority areas for management.

Future work should capture macaque ranging more comprehensively. Continuous GPS telemetry and related methods would allow seasonal and time-of-day variation in macaque occurrence to be described in greater detail. Such information would support more precise decisions about when, where, and along which trail sections management resources for hazing, patrols, and visitor warnings should be concentrated.

In parallel with this study, a web-based reporting form is being prepared to record animal encounters, physical contact, and rule violations. Once operational, this system could provide more direct information on where and how often conflicts between wildlife and visitors occur. These data would allow encounter potential to be evaluated against observed conflict locations and would also support updates to management plans and priority zones.

In parallel with this study, local management bodies, including the Natural Parks Foundation and the Ministry of the Environment, have been preparing a non-public Japanese macaque management manual for park managers, with expert input from the authors. Focused hazing is also being implemented in priority management areas. Future research should evaluate over the long term whether these interventions reduce excessive proximity between visitors and macaques and decrease rule violations. The framework proposed here can therefore serve not only to identify priority sections, but also to provide a baseline for evaluating subsequent management interventions.

## 5. Conclusion

This study visualized the spatial overlap between park visitors and Japanese macaques in Kamikochi as a relative index of encounter potential. Priority sections were concentrated in the central part of the study area and showed especially high values in the frontcountry, corresponding to the strolling area in the zoning scheme. This indicates that the relative potential for encounters between visitors and macaques is not evenly distributed across the study area but is concentrated in a limited part of the landscape.

The main contribution of this study is that it treats overlap itself, rather than visitor counts alone, as a management indicator. Protected-area recreation research has often documented wildlife avoidance, displacement, or temporal adjustment in response to human use. This study complements that perspective by focusing on a situation in which wildlife does not necessarily avoid people and visitor use and wildlife space use continue to overlap. Because the Kamikochi frontcountry is important for both visitors and macaques, management should not rely solely on the expectation that wildlife will spatially separate from areas of high human use. Instead, it is necessary to identify where management efforts should be concentrated within shared-use space.

This perspective can support the implementation and evaluation of management measures in Kamikochi and may also provide a practical framework for spatial prioritization at other sites that pursue wildlife watching and ecotourism without relying on provisioning.

### Management implications

This study provides a spatial basis for prioritizing visitor management in Kamikochi, where visitors and Japanese macaques share the same frontcountry landscape. Because high encounter potential was concentrated in the central frontcountry, limited resources for patrols, visitor warnings, wildlife-viewing management, and hazing should be focused on sections where visitor use and macaque space use most strongly overlap. The results also show that visitor counts alone are insufficient: areas with many visitors or many macaques did not always correspond to high encounter potential when the two distributions were spatially offset. The proposed framework helps translate broad-scale human mobility data into trail-level management decisions and can be applied before and after interventions to evaluate whether targeted actions reduce close approaches, rule violations, or other problematic encounters. It may also inform non-provisioning wildlife-watching sites where wildlife does not consistently avoid visitors.

## Acknowledgements

We are deeply grateful to the staff of the Kamikochi Branch of the Natural Parks Foundation for their extensive support during the fieldwork. We also thank the Ministry of the Environment, Nagano Prefecture, and Matsumoto City for granting the necessary permissions for this study. We are grateful to the local businesses and operators in Kamikochi for kindly accepting and supporting our research activities. We also thank the staff of the Center for the Study of Co*Design and the Cross-Boundary Innovation Program at The University of Osaka for their valuable advice in conducting this interdisciplinary study. This work was supported by the Collaboration Research Program of IDEAS, Chubu University (Grant Numbers IDEAS202402 and IDEAS202501 to YO); MEXT/JSPS KAKENHI (Grant Number JP22K18052 to YO); MEXT/JSPS KAKENHI (Grant Numbers JP26K02099 and JP24H00116 to TM); a research grant from the Institute of Mountain Science, Shinshu University (2026-1501 to TM); and the River Fund of the River Foundation (2026-5211-039 to TM).

